# The structural dynamics and molecular coupling in the slow inactivation of a prokaryotic voltage-gated sodium channel

**DOI:** 10.1101/2025.08.14.670348

**Authors:** Katsumasa Irie, Shuo Han, Sarah Applewhite, Yuki K. Maeda, Joshua Vance, Shizhen Wang

**Affiliations:** Department of Biophysical Chemistry, School of Pharmaceutical Science, Wakayama Medical University, Wakayama, 640-8156, Japan; Division of Biological and Biomedical Systems, School of Science and Engineering, University of Missouri-Kansas City, Kansas City, MO 64110 USA

## Abstract

Voltage-gated sodium (Nav) channels initiate and propagate action potentials in many excitable cells. Upon repetitive activation, the conductance of Nav channels gradually decreases on a timescale ranging from seconds to minutes, a phenomenon known as slow inactivation, which is crucial for regulating the excitability of many cells. Many studies indicated that slow inactivation is associated with conformational changes at the selectivity filter, but the underlying mechanisms remain unclear. By examining the conformational dynamics of a prokaryotic NavAb channel using single-molecule FRET (smFRET), our work revealed the transitions of its selectivity filter among three distinct conformational states and showed that activating voltages enriched the high-FRET conformations, potentially associated with slow inactivation. We further identified L176 in the selectivity filter P1 helix and T206 in the pore-forming S6 helix as residues coupling the primary and slow inactivation gates by showing that the additional L176F mutation stabilizes the S6 C-terminal deletion opening mutant in the closed state. Consistently, our smFRET results also indicated that the high FRET conformation of the selectivity filter was markedly attenuated in the S6 C-terminal deletion opening mutant, but reverted by the L176F mutation. Open-pore blocker lidocaine has been shown to prevent eukaryotic Nav channels from entering the slow inactivation state. Moreover, our smFRET studies showed that it diminished the high FRET conformation of the NavAb selectivity filter in a dose-dependent manner, while the L176F mutation, again, markedly reversed the lidocaine effects. Collectively, our studies suggested that slow inactivation in the NavAb channel results from the collapse of the selectivity filter pore, as revealed by the high FRET conformation in our smFRET measurements. The L176 in the selectivity filter and T206 in the pore-forming S6 helix coordinate conformational changes of the slow inactivation gate at the selectivity filter and the primary gate at the helix bundle crossing, providing the structural basis for slow inactivation in prokaryotic voltage-gated sodium channels.

## Introduction

Voltage-gated sodium channels (Navs) mediate Na^+^ diffusion across cell membranes to initiate action potentials, which are essential for nerve firing or the contraction of skeletal or cardiac muscle ^1,2^. Dysfunction in Nav causes numerous channelopathies, including epilepsy, chronic pain syndromes and cardiac arrhythmias ^3,4^. Immediately after activation, Navs quickly enter the fast inactivation state, allowing the cell membrane potential to be repolarized and initiating another action potential ^5^. The fast inactivation results from voltage-dependent blocking of Na^+^ permeation pores by the linker carrying the ‘IFM’ motif, known as the fast inactivation gate, between domains III and IV of Nav channels ^5,6^. Nav channels also undergo a distinct slow inactivation, which occurs over seconds to minutes, serving as a critical mechanism to prevent persistent nerve firings or muscle contractions ^7^.

Prokaryotic Nav channels share highly conserved structures with eukaryotic orthologs ^8,9^. Due to their simpler structures and amenability to crystallographic and cryo-EM analyses, studies on them have provided invaluable mechanistic insights into the ion selectivity, voltage gating and inactivation of Nav channels ^10–13^. Prokaryotic Nav channels lack fast inactivation gates and, therefore, do not exhibit the fast inactivation typically seen in eukaryotic Nav channels, but their conductance to sodium, upon repetitive activation, does gradually decrease in the time scale of seconds to minutes, a behavior similar to the slow inactivation in eukaryotic Nav channels ^9,14,15^. Indeed, electrophysiological analyses suggested that the collapse of the selectivity filter pore perhaps serves as a shared structural mechanism for slow inactivation in both prokaryotic and eukaryotic Nav channels ^12,15–17^. However, the underlying mechanisms remain elusive, particularly the lack of direct measurements of conformational changes at the selectivity filter during Nav channel transitions from conductive to inactivated states.

In the present work, we leverage NavAb as a structural model to uncover the real-time conformational dynamics of the selectivity filter and the changes induced by activating voltages, thereby understanding the molecular mechanisms driving slow inactivation. We performed smFRET measurements on NavAb channels reconstituted into liposomes and showed that the selectivity filter spontaneously transited among 3 distinct conformational states, with activating voltages enriching the high FRET conformation. It has been previously established that S6 helix C-terminal deletion and lidocaine attenuate slow inactivation; our results indicate that they also diminish the high FRET conformations. Moreover, we identified Leu176 as the critical residue coupling the primary and slow inactivation gates. We showed that the introduction of an additional L176F mutation in the S6 C-terminal deletion mutant, previously crystallized in the opening state, enabled it to adopt a closed conformation in the crystal structure. Our smFRET studies, combined with electrophysiological and crystallographic analyses, consistently indicated that the high FRET, constricted conformation in the selectivity filter represents the slow inactivated conformation, with L176 and T206 mediating conformational talk between the primary gate at bundle crossing and the slow inactivation gate at the selectivity filter.

## Results

### 1. Activating voltages alter the conformational landscapes of the NavAb selectivity filter

So far, the conformational changes associated with slow inactivation in the NavAb channel remain highly speculative, based on crystallographic or cryoEM studies. Thus, we implemented smFRET on NavAb channels reconstituted into liposomes to uncover the structural dynamics underlying slow inactivation. To achieve the studies, we constructed and purified functional tandem dimeric NavAb channel proteins ^18^, which carry cysteine mutations at either E189 or V190 residues at the P2 helix, so that the two cysteines in a tetrameric channel are located at the diagonal subunits (Fig. 1a). The Cy3 and Cy5 FRET fluorophore pair carrying maleimide reactive groups were conjugated to the two cysteine residues, Cy3/Cy5 FRET changes will report the real-time conformational changes of the NavAb selectivity filter. We applied three different electrical potentials (-85, 0, and 120 mV) on NavAb channel proteins in liposomes, generated by intra- and extraliposomal K^+^ gradients in the presence of valinomycin (Fig. 1b). This method has been successfully used in our previous work to examine voltage-dependent conformational changes in voltage-gated proton channels ^19,20^. The wild-type NavAb is inactivated even at deep potentials, but the N49K mutation shifts the activation curve by 75 mV to a half-activation voltage of -22 mV ^10,14^. In this study, the N49K background mutation (named BG in all figures if no additional mutations) was introduced into all mutants, so they were in deactivated, partially activated, or fully activated states at -85, 0, or 120 mV, respectively. Importantly, tandem dimeric NavAb constructs carrying the V190C mutation expressed in SF-9 cells produced functional channels (Extended Fig. 1a and b), and purified tandem dimeric NavAb proteins were also functional, as reported in our previous study ^18^. Our smFRET measurements clearly showed that FRET between Cy3/Cy5 fluorophores labeling at V190C residues at the diagonal subunits exhibits spontaneous transitions among 3 distinct states, suggesting that the NavAb selectivity filter pores may undergo remarkable conformational changes (Fig 1c, extended Fig 2a). Importantly, similar changes were also observed by switching the fluorophore labeling sites to the adjacent E189C (Fig. 1c and extended Fig. 3a). FRET histograms from both the E189C and V190C labeling sites are fitted well with 3 Gaussian components, which resulted in over 99% decreases in the sum of squared errors (Fig 1d). The 3 Gaussian peak centers determined from the E189C labeling sites are slightly higher than those from the V190C sites, matching well with the fact that diagonal E189C residues are closer than diagonal V190C residues (Fig. 1a, e). The high FRET peaks revealed by our smFRET measurements are very close to those predicted from NavAb structures ^11^, and FRET changes between high and medium FRET peaks also match well with those predicted by cryo-EM structures of a prokaryotic NavEh channel at presumptive conductive and slow-inactivated states ^12^. All these data suggest that our smFRET measurements accurately report the structural changes in the NavAb selectivity filter. To analyze the structural dynamics fairly with smFRET data from both E189C and V190C labeling sites together, we used a 3-state kinetic model with FRET peak centers fixed at 0.25, 0.55, and 0.8, named as low (L), medium (M), and high (H) FRET states (Fig. 1f). From both labeling sites, our smFRET contour maps and histograms indicated that the high FRET conformations were significantly enriched at activating voltages, so it is reasonable to assume that they may represent the NavAb selectivity filter in the inactivated state (Fig. 1g and h).

**Figure 1.**
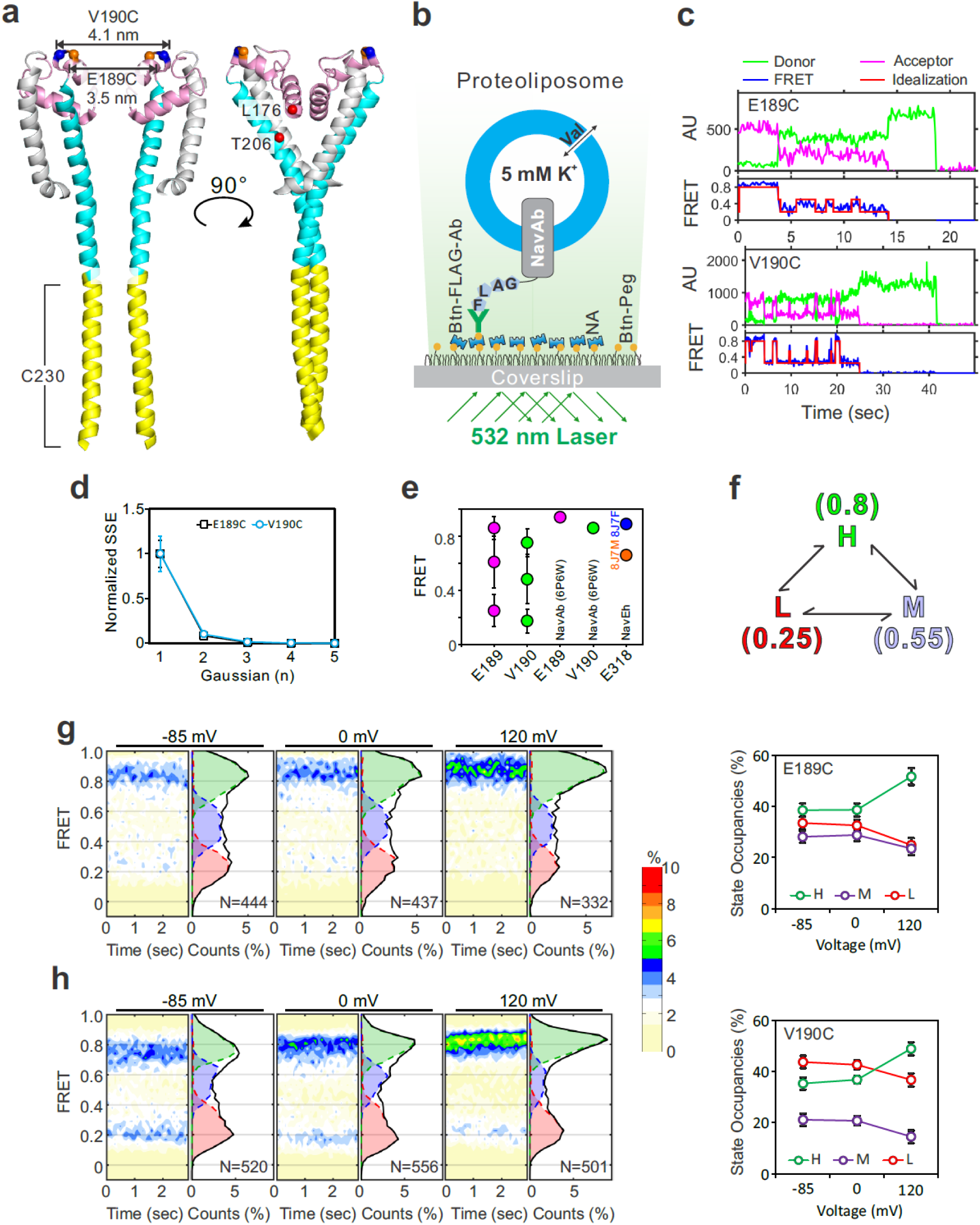
Activating voltage promotes the constricted conformation at the NavAb selectivity filter. **a**. Cartoon of the NavAb pore domain, with 2 subunits removed for clarity (PDB: 5VB2). The pore-forming helices are colored cyan, and the P1/P2 helices of the selectivity filter are colored pink. The C-terminal helices deleted in the ΔC230 mutant are colored yellow, and the L176 and T206 residues mutated are highlighted as red spheres. The Cα carbon atoms of E189C and V190C, at the diagonal subunits for fluorophore labeling, were highlighted by orange and blue spheres, respectively. **b**. The experimental setup for examining the conformational dynamics of the NavAb selectivity filter using smFRET. The fluorophore-labeled NavAb channels were reconstituted into liposomes (POPE/POPG=31, w/w), and the proteoliposomes were immobilized on the PEGylated surface through neutravidin and biotinylated anti-histag or anti-FLAG tag antibodies, so only those with the C-terminus of the NavAb channel facing outside were retained for smFRET imaging. Transliposomal electrical potentials were generated by transliposomal K^+^ gradients and valinomycin, with the intraliposomal K^+^ concentration as 5 mM and extraliposomal K^+^ concentration as 150, 5, and 0.04 mM for -85, 0, and 120 mV, respectively. **c**. smFRET traces showed that FRET between Cy3/Cy5 fluorophores conjugated at E189C or V190C at the P2 helix in the NavAb selectivity filter exhibits transitions among 3 distinctive FRET states. **d**. smFRET histograms of E189C and V190C labeling sites were fit with 1-5 Gaussian components. Three Gaussians reduced the sum of squared errors (SSE) by >99% and were therefore used for all subsequent state occupancy and kinetic analyses. **e**. The centers of FRET states from Gaussian fitting of smFRET histograms from the diagonal E189C and V190C labeling sites. The FRET efficiencies of the diagonal E189 and V190 predicted from structures of NavAb (6P6W) or diagonal E318 from the structures of NavEh (8J7F and 8J7M) were also included for comparison. **f**. The kinetic model for analyzing smFRET traces collected from E189C and V190C labeling sites. To calculate and compare FRET state occupancies fairly, smFRET traces collected from E189C and V190C sites were idealized into low, medium, and high FRET states, with peak centers fixed at 0.25, 0.55, and 0.8, respectively. **g-h**. Contour maps and histograms of smFRET traces collected from E189C (**g**) and V190C (**h**) labeling sites indicated that strong activating voltages enriched the high FRET populations. The FRET state occupancy data (right panels), calculated as fractional areas from histograms with uncertainties reported as standard errors, clearly showed that strong activating voltages enriched the high FRET state but diminished the medium and low FRET states.

### 2. SmFRET measurements revealed conformational changes in the selectivity filter induced by mutations that alter slow inactivation in NavAb channels

The S6 helix is a long, uninterrupted α-helix that affects both activation and slow inactivation of the NavAb channel ^21^. The cytosolic helix bundle formed by four S6 helices of a tetrameric NavAb channel is a structural feature shared among prokaryotic Navs ^22^. Deletion of the S6 helix bundle enhances the early phase of slow inactivation while blocking the late use-dependent inactivation of the NavAb channel (Fig 1a). In addition, mutation of the S6 residue close to the selectivity filter, T206, also markedly alters its slow inactivation behavior ^16,22^. L176 at the NavAb selectivity filter is very close to T206; thus, they may couple the conformational changes between the primary helix bundle crossing gate and the selectivity filter slow-inactivation gate. Indeed, our electrophysiological data indicated that mutations to bulky side chain residues like Phe (L176F) and Trp (L176W) significantly altered the slow inactivation of NavAb channels (Fig. 2a and extended Fig. 4a, b, c). The voltage dependencies of L176F and L176W activation were almost unchanged, while their half-inactivation voltages were shifted negatively by over 30 mV (Table 1). We also observed similar changes in slow inactivation from the mutant carrying a C-terminal deletion to Q230, named ΔC230 (Fig. 2a, extended Fig. 4d, e) ^16^.

**Figure 2.**
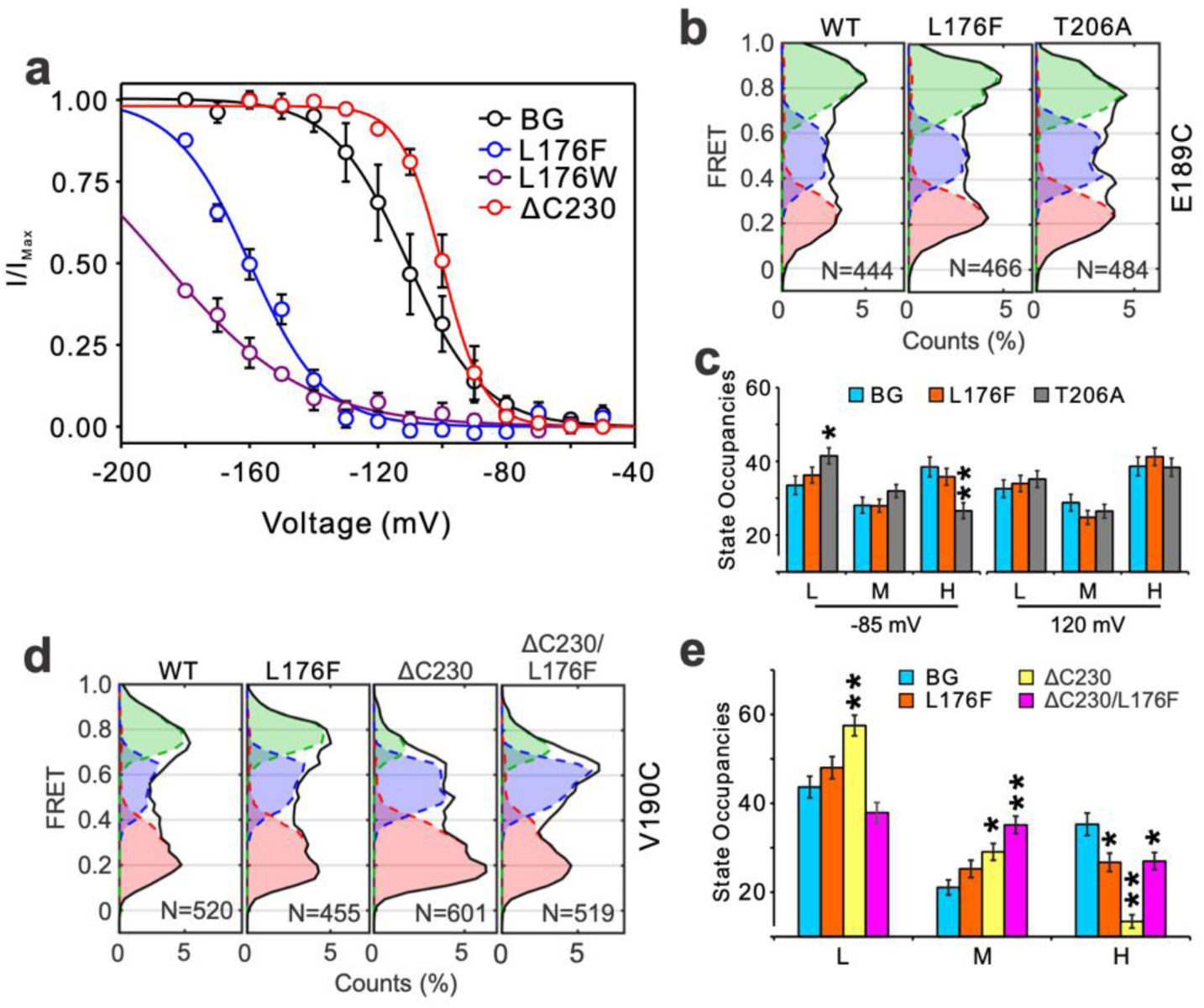
SmFRET measurements revealed conformational changes in the selectivity filter induced by mutations that alter slow inactivation in the NavAb channel. **a**. Steady-state inactivation of NavAb BG, L176F, L176W, and ΔC230 mutants. **b**. The histograms of smFRET traces from E189C labeling sites collected without or with the L176F and T206A background mutations under -85 mV membrane voltages. **c**. FRET state occupancies of smFRET traces from E189C labeling sites without or with the L176F and T206A background mutations under -85 mV and 120 mV membrane voltages. T-tests were performed to examine significance levels relative to BG, with * denoting p<0.05 and ** denoting p<0.001. **d.** The histograms of smFRET traces from V190C labeling sites collected without or with the L176F C-terminal deletion ΔC230 and both background mutations under -85 mV membrane voltages. **e**. FRET state occupancies of smFRET traces from V190C labeling sites without or with the L176F, C-terminal deletion ΔC230 and both background mutations under -85 mV membrane voltages. T-tests were performed to examine significance levels relative to BG, with * denoting p<0.05 and ** denoting p<0.001.

**Table 1.**
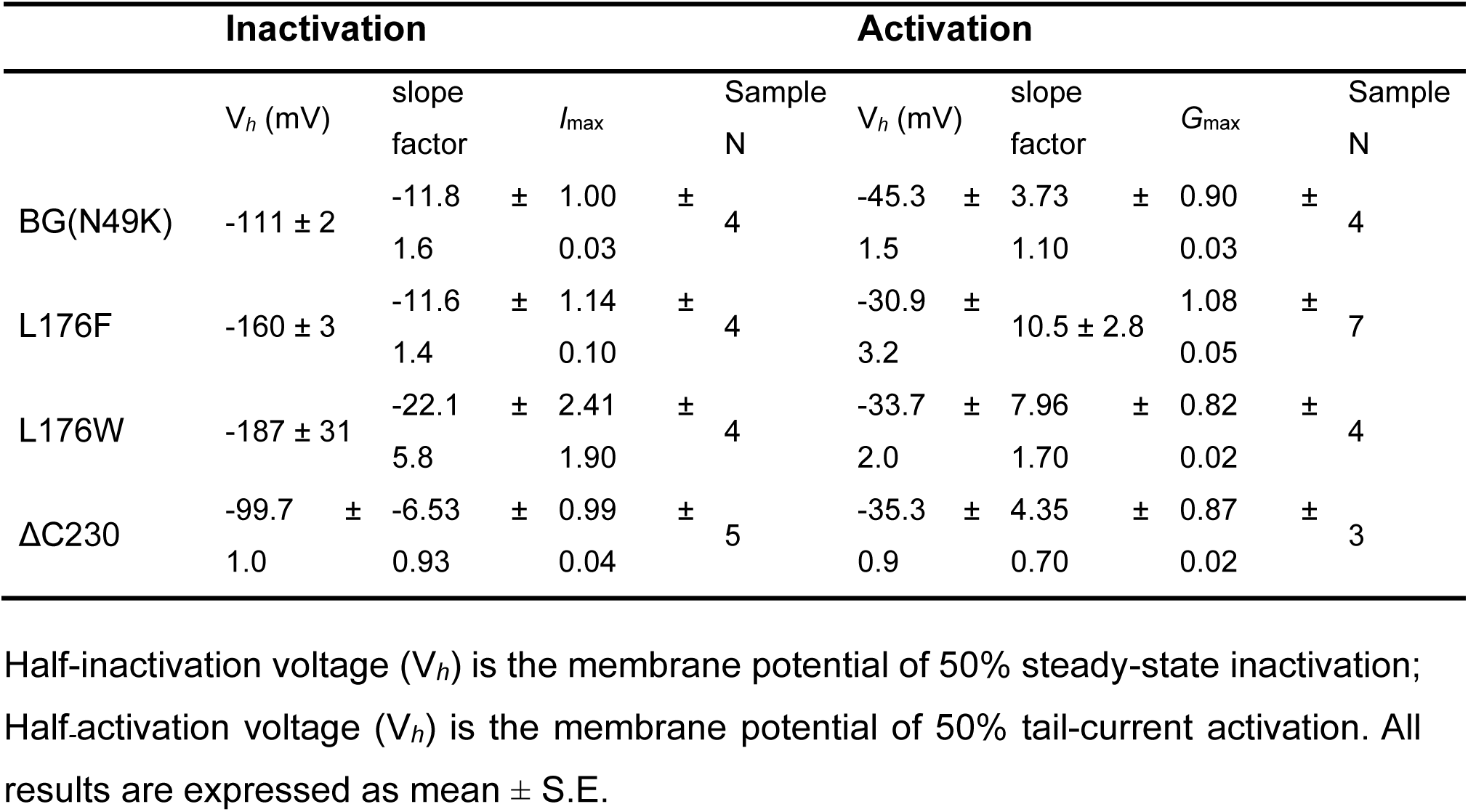
Half-inactivation and activation voltages of NavAb mutants.

We examined the effects of the T206A, L176F, and ΔC230 mutations on the NavAb selectivity filter conformation using smFRET. These tandem dimeric NavAb constructs produced functional channels, except L176W (Extended Fig. 1b-f). Due to the absence of measurable currents in the L176W tandem dimer, this mutant was excluded from FRET analysis. Our results clearly showed that the T206A mutation significantly depletes the high-FRET population at -85 mV, consistent with its functional effects of attenuating slow inactivation in NavAb channels. However, its effects became indistinguishable under strong activating voltages (Fig 2b-c, extended Fig. 3b). Dramatic changes, however, were observed from the ΔC230 mutant, where the high FRET population was almost absent, even under strong activating voltages, in sharp contrast to the T206A mutant (Fig 2d, e, extended Fig 2b). Surprisingly, the L176F mutation does not cause significant changes in the selectivity filter conformation, as indicated by smFRET collected from both the E189C and V190C labeling sites (Fig 2b-e, extended Fig 2b, Fig. 3b). However, it strongly attenuates the ΔC230 effects by enriching the high-FRET state and diminishing the low-FRET state, consistent with its role in promoting slow inactivation (Fig 2d, e). Collectively, our smFRET data from NavAb mutants supported the idea that conformational changes revealed by smFRET measurements are associated with slow inactivation, and that high FRET conformations may underlie slow inactivation in the NavAb selectivity filter.

### 3. T206 and L176 mediate conformational talk between the slow inactivation and primary gates

To reveal the conformational changes associated with slow inactivation, we further obtained crystal structures of the NavAb channel carrying the N49K background mutation (named BG hereafter) and those containing additional mutations, including L176F, ΔC230, and T206A (Extended Tables 1 and 2). Structural alignments indicated that, despite changes in their slow inactivation behaviors, the selectivity filters of L176F, T206A, and ΔC230 mutants exhibit no visible changes (Fig 3a and extended Fig 5). The overall RMSD of all structures before the S6 helix is below 1Å, indicating there is very minimal overall structural change in all mutants (Fig 3b).

**Figure 3.**
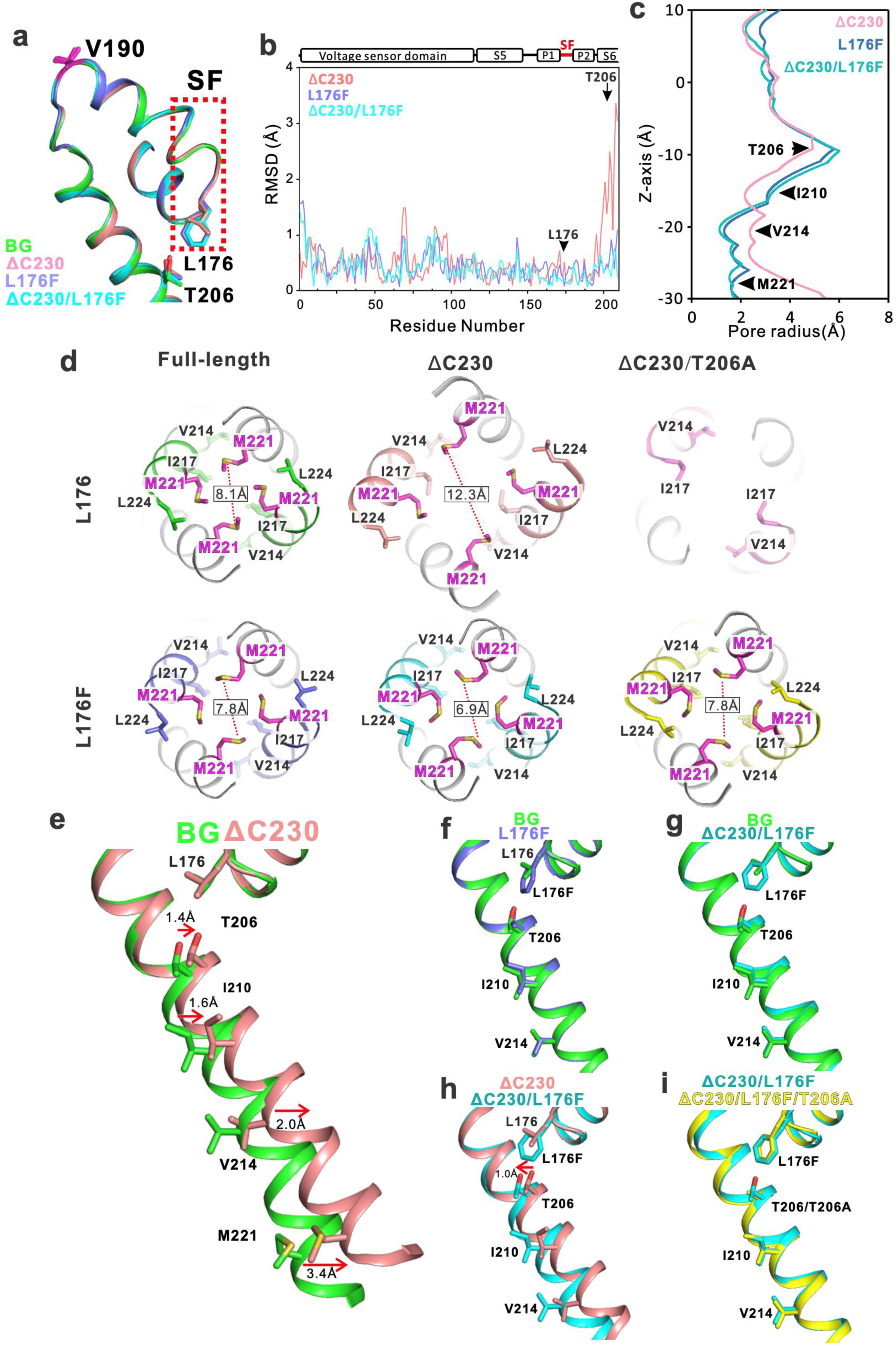
The molecular coupler mediates conformational changes between the selectivity filter and the S6 helix bundle crossing gate. **a**. Superimposition of the selectivity filter of crystal structures of the BG, L176F, ΔC230 and ΔC230/L176F mutants. The T206 and L176 residues, which couple the selectivity filter to the S6 helix, and V190 for fluorophore labeling, are highlighted as sticks. **b**. Root Mean Square Deviation (RMSD) of pore domain residues of mutant channels, L176F, ΔC230 and ΔC230/L176F, against NavAb BG mutant carrying only the N49K background mutation. **c**. Pore diameter profiles of NavAb BG, L176F, ΔC230 and ΔC230/L176F mutants. **d**. Bottom views of the helix bundle cross gates of NavAb channels without (BG) or with the L176F, ΔC230, ΔC230/L176F and ΔC230/T206A background mutations. They clearly indicated that the L176F mutation stabilizes the ΔC230 opening mutant at the closed state. **e-i**. Superimposition of the selectivity filter and the S6 helix of BG and ΔC230 (**e**), BG and L176F (**f**), BG and ΔC230/L176F (**g**), ΔC230 and ΔC230/L176F (**h**), ΔC230/L176F and ΔC230/L176F/T206A (**i**), viewing from figure1a right direction. The L176F mutation pushes the S6 helix toward the closed state conformation.

However, the RMSD of the ΔC230 mutant around the S6 helix was greater than 1 Å, suggesting a significant difference in the primary gate conformation (Fig 3b). The primary gate of the ΔC230 mutant is very different from that of the full length channel (BG), apparently at the opening state or transiting towards the opening state (Fig 3c and d). Interestingly, the ΔC230/L176F mutant was crystallized in the closed state, indicating that the L176F mutation stabilizes the primary bundle crossing gate at the closed state (Fig 3c, d). In contrast, adding the T206A mutation to the ΔC230 mutant did not result in closure of the primary gate. Rather, the S6 C-terminus forming the primary gate is too flexible to be resolved in the crystal structure (Fig 3d and extended Fig 6). T206 is one of the residues where the pore radius undergoes significant narrowing in the ΔC230 mutant compared to the full-length channel (Fig 3c). The T206A mutation would create a gap that increases the mobility of the S6 helix, resulting in the disappearance of electron density at the S6 C-terminus in the ΔC230/T206A mutant. Our previous studies also showed that the T206A mutation tends to decrease structural resolution ^23^, a finding that can now be attributed to this high mobility of the S6 helix.

Notably, the L176F effects on the primary gate, in fact, were also observable in the crystal structure of the ΔC230/L176F/T206A mutant (Fig 3d). In all these structures, it is clear that residues at the 176 position were very close to T206 in the S6 helix (Fig 3e-i, extended Fig 5, Fig 6). In the ΔC230 mutant, T206 moves toward L176 (Fig. 3e), whereas in the L176F and ΔC230/L176F structures, T206 superimposes well with the BG structure (Fig 3f, g), rather than ΔC230 mutant (Fig. 3h). Their proximity to L176 also exhibits no difference, except for the ΔC230 mutant, as indicated by the distances from the T206 Cβ carbon to the L176 Cγ, Cβ, and Cα carbons (Table 2). Therefore, in the ΔC230/L176F structure, the T206 is pushed away from L176 by more than 1Å through steric hindrance with the T206, which repositions the S6 helix and closes the primary helix bundle crossing gate (Fig. 3e, h). A similar effect was observed in the ΔC230/L176F/T206A triple mutant (Fig. 3i). A similar, even stronger pushing effect was observed with L176W (Table 2), consistent with our electrophysiological data (Fig. 2a). However, the increased B-factor of the crystal structure and our inability to measure the L176W tandem dimer suggest that excessive steric hindrance may destabilize the overall channel structure.

**Table 2.** Root mean square deviation (RMSD) and distance between the 176 residues and T206 of the NavAb channels.

| | RMSD with | | Distance from T206 Cβ to | | | |
| --- | --- | --- | --- | --- | --- | --- |
| | BG mutant (Å) | | Nearest atom | Cγ | Cβ | Cα |
|  | 176 Residues | Thr206 | of 176 residues (Å) |  |  |  |
| BG | - | - | 6.6 | 6.9 | 8.4 | 9.0 |
| L176F | 0.30 | 0.68 | 4.1 | 6.8 | 8.3 | 8.9 |
| ΔC230 | 0.23 | 2.31 | 6.2 | 6.2 | 7.6 | 8.1 |
| ΔC230/L176F | 0.15 | 0.50 | 4.3 | 6.8 | 8.3 | 8.7 |
| T206A | 0.33 | 0.28 | 6.8 | 7.0 | 8.5 | 9.2 |
| L176W | 0.70 | 0.79 | 4.8 | 7.8 | 9.2 | 9.9 |

Intriguingly, our smFRET data showed that introducing the L176F mutation reversed the conformational changes at the selectivity filter induced by the ΔC230 mutation, increasing the medium and high FRET populations (Fig. 2d-e). Our crystallographic analyses and smFRET results above clearly demonstrate strong conformational coupling between the primary helix bundle crossing gate and the selectivity filter inactivation gate. It also implied that the slow-inactivation conformation of the selectivity filter could be highly dynamic or unstable compared to that of channels with opening primary gates.

### 4. Open-channel blocker lidocaine alters the NavAb selectivity filter dynamics in the same way as the C-terminal deletion open mutation

As a classical open-channel blocker of eukaryotic Nav channels, lidocaine was found to prevent Nav1.5 and Nav1.7 channels from entering the slow inactivation state ^24,25^. Lidocaine was also shown to block prokaryotic NavAb channels with high affinities at the inner cavity immediately underneath the selectivity filter pore ^26^. In the present work, we examined the effects of lidocaine on the selectivity filter conformation using smFRET. Our results indicate that lidocaine depleted the high-FRET conformations in a clear concentration-dependent manner, reaching saturation at 0.5 mM (Fig. 4a, b). The observed effects of lidocaine on NavAb selectivity filter conformation matched well with its functional results of preventing Nav channels from entering the slow inactivation state, supporting that high FRET conformation represents the selectivity filter at the inactivated state. Interestingly, introducing the L176F mutation restored the high FRET population, even in the presence of 5 mM lidocaine, an effect similar to that observed from the ΔC230 mutant. (Fig. 4a-b). By analyzing transition kinetics among L, M, and H FRET states using FRET state occupancy data, our smFRET data indicated that lidocaine promotes transitions from L to H and M to H FRET states, with a half-effective concentration close to 0.05 mM, very close to the apparent Kd of 0.1 mM reported by by electrophysiological studies ^26^ (Fig. 4c). The transition kinetics data collectively suggested that high-FRET conformation underlies the slow inactivation of the NavAb channel, while the medium FRET could be the early phase of slow inactivation.

**Figure 4.**
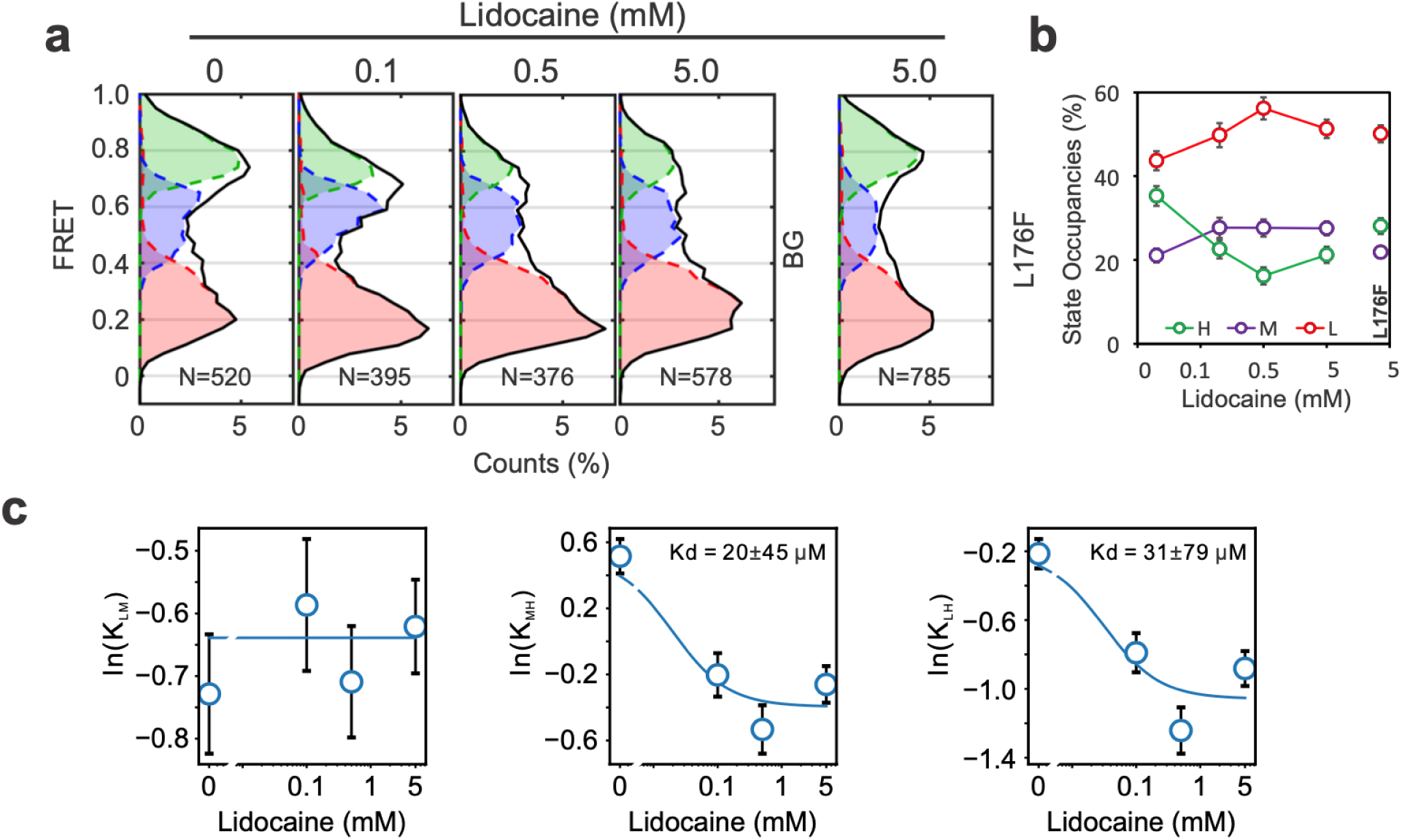
Lidocaine modifies the conformation of the NavAb selectivity filter in a dose-dependent manner. **a** and **b**. FRET histograms (**a**), state occupancies of smFRET data collected from V190C labeling sites, without (BG) or with the L176F background mutation, in the presence of different concentrations of lidocaine under 0 mV. Lidocaine depleted the high FRET population in a dose-dependent manner, while the L176F mutation, which enhances slow inactivation, restored the high FRET conformation of the selectivity filter. **c**. Changes in equilibrium constants for transitions between L and M (K_­_), M and H (K_­_), and L and H (K_­_) FRET states with lidocaine concentration were fitted to the Hill equation with a Hill coefficient of n=1. The uncertainties of the equilibrium constant K were reported as standard errors. Only transition equilibrium constants between M and H and L and H states are changed by lidocaine, with an apparent Kd very close to 0.1 mM, close to that determined by electrophysiological studies.

## Discussions

Slow inactivation is a key mechanism of Navs for regulating the frequencies of nerve firings and muscle contractions ^7^. Residues in the selectivity filter, pore, and voltage-sensor domains of Navs were found to affect slow inactivation, but many studies have suggested that slow inactivation involves the collapse of the selectivity filter pore, a mechanism quite similar to C-type inactivation in potassium channels ^27,28^. The prokaryotic NavAb channel also exhibits slow inactivation behaviors ^9,16^, but the underlying structural mechanisms remain unclear. In the present work, we conjugated Cy3/Cy5 FRET to the diagonal E189C or V190C residues in the selectivity filter at the P2 helix to examine conformational dynamics in the NavAb selectivity filter. Our smFRET results indicated that the NavAb selectivity filter is very different from that of the potassium channel, which exhibits very rare FRET transitions, suggesting a fundamental difference in their structural basis of ion selectivity ^29^. SmFRET data from both E189C and V190C labeling sites consistently showed 3 distinct FRET states. The high FRET populations of its selectivity filter were (a) enriched by strong activating voltages, (b) attenuated by the S6 helix T206A, ΔC230 mutations attenuating the slow inactivation, (c) diminished by the lidocaine, which was found to prevent Nav channels from entering slow inactivation state, and (d) restored by the selectivity filter mutation L176F promoting slow inactivation. We further calculated the equilibrium constants (Kij) for transitions between L and M (LM), L and H (LH), M and H (MH) using FRET state occupancy data. Our results indicated that activating voltages significantly promote transitions from low and medium to high FRET states; T206A and ΔC230 mutations, and lidocaine, promote transitions from high to medium and low FRET states, with the L176F mutation reversing their effects (Fig 5a, b). Our results support the idea that the high FRET constricted selectivity filter conformations may be associated with slow inactivation in the NavAb channel, while low FRET conformations may represent the dilated selectivity filter pore conformation, which is conductive (Fig. 5c). The functional roles of the medium FRET conformation, however, are yet to be established. Since the FRET Cy3/Cy5 fluorophore pair actually reports relative movements between two diagonal P2 helices, the medium FRET state could represent an asymmetric selectivity filter pore, with one subunit adopting the high-FRET inactivated conformation and the other adopting the low-FRET conductive conformation. The selectivity filters of the eukaryotic Nav channels are asymmetrical, so their structural basis of slow inactivation could be even more complicated ^30^.

**Figure 5.**
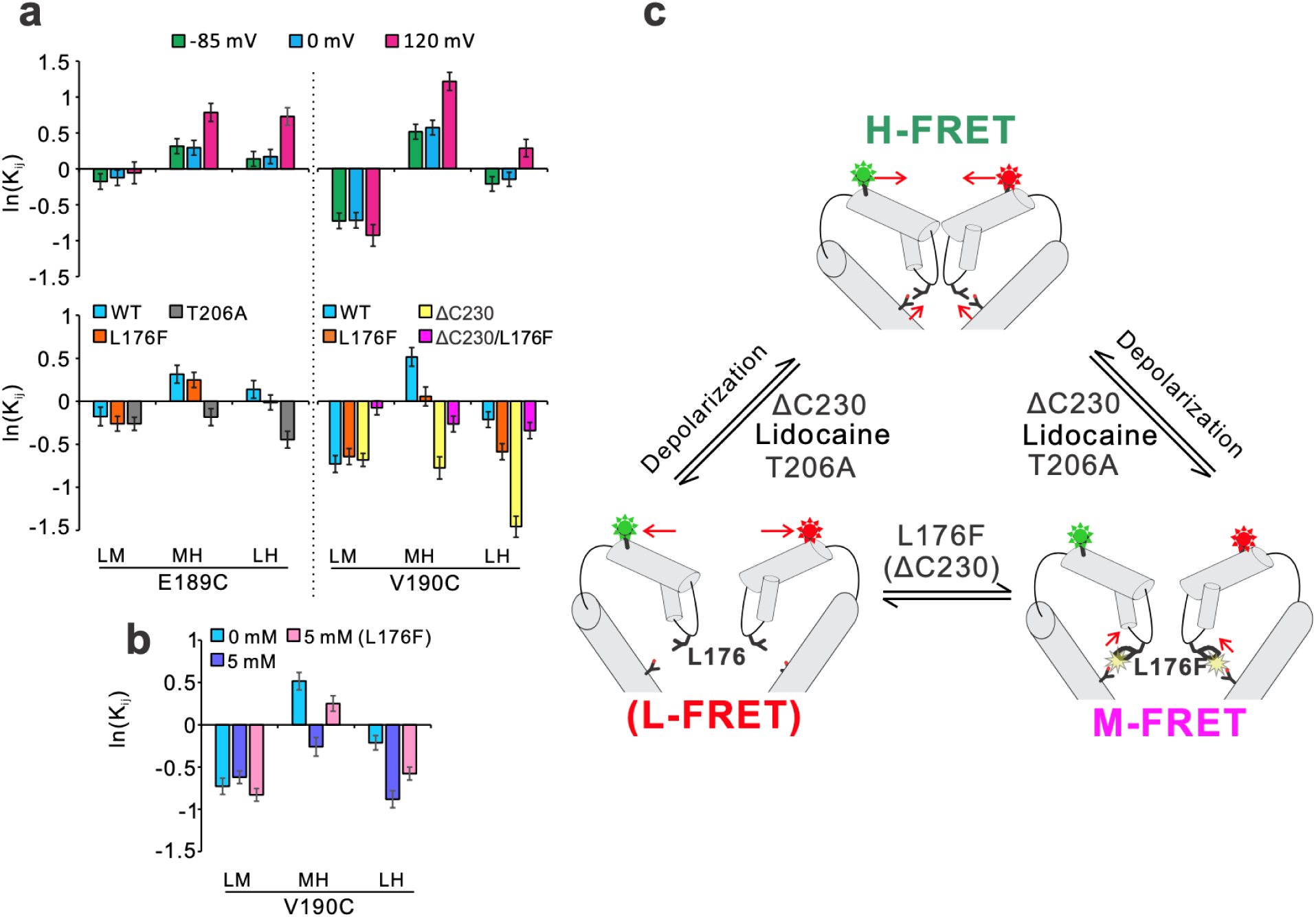
Selectivity filter conformational dynamics underlying slow inactivation in the NavAb channel. **a** and **b**. Effects of voltage (**a,** upper panel), mutations under -85 mV (**a,** lower panel) or lidocaine under 0 mV (**b**) on equilibrium constants Kij of transitions between L and M (LM), M and H (MH), L and H (LH) FRET states. The transition equilibrium constants Kij were calculated from FRET state occupancy data obtained from either the E189C or V190C labeling sites, as described in the materials and methods section. The uncertainties of Kij were reported as standard errors. **c**. Structural basis underlying slow inactivation in the NavAb channel. The low and high FRET populations correspond to the conductive and slow inactivation states, respectively. T206A, lidocaine (lido), and C-terminal deletion (ΔC230) promote transitions from high to medium or low FRET conductive state conformations, while depolarized voltages (depo) promote transitions from low and medium to high FRET inactivated state conformations. Only the pore domains of two subunits were shown, with the front and rear subunits removed for clarity.

NavAb channels, whether open or closed, also exhibit almost identical selectivity filter conformations^11,21^. Only very small changes have been observed in NavAb crystal structures with unusual 2-fold symmetry, but their roles in slow inactivation remain to be established ^13^. So far, no coupled conformational changes were identified between the primary and inactivated gates in all available NavAb structures, a hallmark feature of slow inactivation ^7,17^. One possibility is that the primary gates of all ‘opening’ NavAb structures were not open sufficiently to stabilize the selectivity filter in the slow inactivated conformation. Actually, this is true that in the KcsA potassium channel, in which the selectivity filter does not transit to the inactivated conformation until the primary gate opens by over 20 angstroms ^27,28^. So, to capture the inactivated NavAb selectivity filter, we may need mutants carrying a reinforced coupler, such as L176F, plus mutations that open its primary gate sufficiently wide, such as glutamate mutations at the helix bundle crossing gate in potassium channels ^31,32^. Recent cryo-EM analyses of a homotetrameric NavEh channel, cloned from eukaryotic single-cell algae, revealed remarkable changes at the P2 helix in the selectivity filter that may be associated with slow inactivation ^12^. These NavEh structures revealed a partially unfolded P2 helix, with its selectivity filter adopting a more dilated conformation, which may be stabilized by extracellular loops (ECLs) seen in eukaryotic Navs but absent in the NavAb channel. Nevertheless, if we define the gating status of these NavEh structures based on their primary gates. Then, the open structure with a constricted conformation could correspond to the inactivated selectivity filter, and the predicted FRET efficiency changes match well with our high FRET data (Fig 1e). While this structural model requires further evaluation to be fully confirmed, it provides a plausible explanation for our observations.

Our research has revealed multiple dynamic transitions associated with slow inactivation using smFRET measurements and further identified L176 as a coupling factor between the primary gate and inactivation gate through crystallographic analysis. These discoveries provide crucial insights into the mechanism of slow inactivation, a process that has long remained a challenging phenomenon to resolve. The integration of approaches that address both static and dynamic features of Nav channels will pave the way for a comprehensive understanding of the molecular basis of slow inactivation in eukaryotic Nav channels.

## Materials and Methods

### Site-directed mutagenesis and construction of NavAb mutants

The NavAb mutated DNAs were subcloned into the modified pBiEX-1 vector (Novagen) that was modified by replacing the fragment from the NdeI site to the SalI site in the multi-cloning site with that of the previously described modified pET-15b (Novagen) vector between BamHI and SalI ^33,34^. The polymerase chain reaction accomplished site-directed mutagenesis using PrimeSTAR® Max DNA Polymerase (Takara Bio). All clones were confirmed by DNA sequencing. The plasmids expressing tandem dimeric NavAb proteins were constructed by inserting 2 copies of NavAb cDNAs into the pET28a vector at NdeI-BamHI and BamHI-SalI sites, with a flexible linker sequence encoding double ‘GGGS’ and the FLAG tag ^18^.

### Electrophysiological measurement in insect cells

The recordings were performed using SF-9 cells. SF-9 cells (ATCC#CRL-1711) were grown in Sf-900™ II medium (Gibco) complemented with 1% 100× Antibiotic-Antimycotic (Fujifilm-Wako) at 27°C. Cells were transfected with target channel-cloned pBiEX vectors and enhanced green fluorescent protein (EGFP)-cloned pBiEX vectors using polyethyleneimine (PEI) reagent ^33,34^. PEI was solubilized into distilled water, and the pH of the 1 mg/mL PEI solution was adjusted to 7.0 with NaOH. First, the channel-cloned vector (3.3 μg) was mixed with 1.6 μg of the EGFP-cloned vector in 150 μL of the culture medium. Next, 15 μL 1mg/mL PEI solution was added, and the mixture was incubated for 10 min before the transfection mixture was gently dropped onto cultured cells. After 24-48 h incubation, the cells were used for electrophysiological measurements. Cancellation of the capacitance transients and leak subtraction were performed using a programmed P/10 protocol delivered at -140 mV. The bath solution was changed using the Dynaflow® Resolve system. All experiments were conducted at 25 ± 1°C using a whole cell patch clamp recording mode with a HEKA EPC 10 amplifier and Patch master data acquisition software (v2×73). Data export was done using Igor Pro 9.05 and NeuroMatic (version 3.0b). All sample numbers represent the number of individual cells used for each measurement. Cells with leak currents less than 1 nA were used for measurements. When any outliers were encountered, these outliers were excluded if any abnormalities were found in other measurement environments and were included if no abnormalities were found. All results are presented as mean ± SE. The graph data were plotted using SigmaPlot 14.

### Protein expression and purification

Proteins were expressed in the Escherichia coli KRX strain (Promega). Cells were grown at 37°C to an OD_­_of 0.6, induced with 0.1% rhamnose (Fujifilm-Wako), and grown for 16 h at 20°C. The cells were suspended in TBS buffer (20 mM Tris-HCl pH 8.0, 150 mM NaCl) and lysed using LAB1000 (SMT Co., LTD.) at 12,000 psi. Low-speed centrifugation removed cell debris (12,000×g, 30 min, 4°C). Membranes were collected by centrifugation (100,000×g, 1 h, 4°C) and solubilized by homogenization in TBS buffer containing 30 mM n-dodecyl-β-D-maltoside (DDM, DOJINDO). After centrifugation (40,000×g, 30 min, 4°C), the supernatant was loaded onto a HIS-Select® Cobalt Affinity Gel column (Sigma). The protein bound to the cobalt affinity column was washed with 10 mM imidazole in TBS buffer containing 0.05% lauryl maltose neopentyl glycol (LMNG, Anatrace) instead of DDM. After washing, the protein was eluted with 300 mM imidazole, and the His tag of the protein samples for crystallization was removed by thrombin digestion (overnight, 4°C). Eluted protein was purified on a TSKgel G4000SWXL (TOSO Corp.) in TBS buffer containing 0.05% LMNG.

### Crystallization and structural determination

Before crystallization, the purified protein was concentrated to ∼10mg ml-1 and reconstituted into a bicelle solution containing a 10% bicelle mixture at 2.8:1 (1,2-dimyristoyl-sn-glycero-3-phosphorylcholine [DMPC, Anatrace]: 3-[(3-cholamidopropyl) dimethylammonio]-2-hydroxy propane sulfonate [CHAPSO, DOJINDO]). The 10mg/ml protein solution of NavAb and 10% bicelle were mixed in a 20:1 ratio. Prepared proteins of the full-length NavAb mutants were crystallized by sitting-drop vapor diffusion at 20°C by mixing 300-nl volumes of the protein solution (8–10 mg/ml) and the reservoir solution (9%–11% polyethylene glycol monomethyl ether [PEG MME] 2000, 100 mM sodium chloride, 100 mM magnesium nitrate, 25mM cadmium chloride and 100 mM Tris-HCl, pH 8.4) with mosquito LCP (STP Labtech). The crystals were grown for 1 to 3 weeks, and after growing, the crystals were transferred into the reservoir solution without cadmium nitrate, which was replaced with the following cryoprotectant solutions. The cryoprotectant solution contains 11% PEG MME 2000, 100 mM Tris-HCl pH 8.4, 2.5 M sodium chloride, 100 mM magnesium nitrate, 0.05% LMNG, and 20% (v/v) DMSO.

Prepared proteins of C-terminal deletion NavAb mutants were crystallized by sitting-drop vapor diffusion at 20°C by mixing 300-nl volumes of the protein solution (8–10 mg/ml) and the reservoir solution (7%–9% PEG 6000, 100 mM sodium chloride, 100 mM magnesium nitrate, 100mM cadmium chloride, 10mM copper chloride and 100 mM Tris-HCl, pH 8.4) with mosquito LCP (STP Labtech). The crystals were grown for 1 to 3 weeks, and after growing, the crystals were transferred into the reservoir solution without cadmium nitrate, which was replaced with the following cryoprotectant solutions. The cryoprotectant solution contains 11% PEG 6000, 100 mM Tris-HCl pH 8.4, 2.5 M sodium chloride, 100 mM magnesium nitrate, 0.05% LMNG, and 20% (v/v) DMSO.

All crystallography data were automatically collected at BL41XU and BL45XU of SPring-8 and merged using the ZOO system ^35^. The data were processed by the KAMO system ^36^ with XDS (version 20220110) ^37^. The data sets of the NavAb BG were obtained from a single crystal. The data sets of the L176F and L176W mutants of NavAb were obtained from multiple crystals. Analyses of the data with the STARANISO server (Global Phasing Ltd) ^38^ revealed severely anisotropic crystals. Therefore, the data sets were ellipsoidally truncated and rescaled to minimize the inclusion of poor diffraction data.

A molecular replacement method with PHASER ^39^ provided the initial phase using the structure of NavAb N49K mutant (pdb code: 5yuc) as the initial model. The final model was constructed in COOT (version 0.9.2) ^40^ and refined in Phenix (version 1.18) ^41^. CCP4 package (version 7.0.078) ^42^ was used for Structural analysis. Supplementary Table 1 summarizes data collection and refinement statistics for all crystals. All figures in the present paper were prepared using the program PyMOL 2.5.4 (Schrödinger, LLC, 2015:The PyMOL Molecular Graphics System, Version 1.8). The pore radii were calculated using the HOLE program ^43^.

### Fluorophore labeling, liposome reconstitution, smFRET imaging and statistics

NavAb proteins carrying either E189C or V190C mutations were conjugated with photostable Cy3/Cy5 c5 maleimide (1:1) as described previously ^18,44^. The fluorophore-labeled NavAb proteins were then reconstituted into liposomes containing POPE/POPG (3:1, w/w, Avanti Polar Lipids Inc.) with a protein lipid ratio of 1:4,000 (w/w) using Bio-beads SM2 (Bioad Inc.) to remove detergents, and the NavAb proteoliposomes were either immediately used or stored in a -80 °C freezer. Immediately before smFRET imaging, the NavAb proteoliposomes were extruded through a 200 nm polycarbonate membrane >30 times to obtain unilamellar, uniform-size liposomes, then immobilized on PEG/biotin-PEG (98:2, w/w) passivated coverslip surfaces for imaging through biotinylated anti-Histag or anti-FLAG antibodies (ThermoFisher, MA1-21315-BTIN, MA1-91878-BTIN) ^45^. SmFRET imaging was performed with an objective-based TIRF microscope, and liposomal electrical voltages were generated by transliposomal K^+­^gradients (K^+^_­_/K^+^_­_, mM, 5/150 for -85 mV, 5/5 for 0 mV and 5/0.04 for 120 mV) in the presence of 0.45 µM valinomycin ^20^. SmFRET movies were collected by an exciting laser of 532 nm (∼1.0 W/cm^2^) with a time resolution of 100 ms. The PCD/PCA oxygen scavenger system and ∼3 mM Trolox were included in the imaging buffer to enhance the photostability of the fluorophores ^46^. Raw smFRET movies without any corrections were processed using the SPARTAN software with a point spread function (PSF) window size of 7 pixels. SmFRET traces were then selected using the Autotrace function and further manually inspected according to the criteria described previously ^20,47^. After correction of the crosstalk and gamma effect, smFRET traces were idealized using the Maximum Point Likelihood algorithm built into the SPARTAN software with a 4 FRET state model, including the low, medium, high FRET states and one additional zero FRET state accounting for the blinking and bleaching FRET events ^48^.

FRET state occupancies was calculated as fractional areas from histograms of all idealized smFRET traces for each experimental condition, with uncertainties reported as standard errors. Steady-state occupancies of the three conformational states (L, M, H) were obtained from FRET histograms and expressed as fractional populations *p*_*L*_, *p*_*M*_, and *p*_*H*_. Pairwise equilibrium constants between states were calculated directly from the occupancies as:

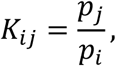

and converted to an energetic representation using:

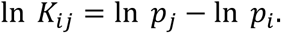

Uncertainties in ln *K* were estimated by propagation of errors from the measured standard errors of the state occupancies. Thermodynamic consistency was verified by confirming closure of the three-state cycle:

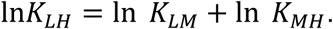

ln *K* values were plotted as a function of lidocaine concentration and fitted with the Hill equation with Hill coefficient n=1:

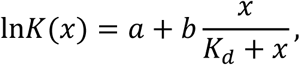

where *a* is the baseline equilibrium, *b* is the ligand-induced change in ln *K*, and *K*_*d*_ is the apparent dissociation constant. Curve fitting was performed by nonlinear least-squares regression to obtain the apparent *K*_*d*_. Data are reported as mean ± propagated standard error.

### Data Availability

The structural data generated in this study have been deposited in the Protein Data Bank under accession codes 9VDQ[https://doi.org/10.2210/pdb9VDQ/pdb]: Crystal structure of voltage-gated sodium channel NavAb N49K mutant, 9VDR[https://doi.org/10.2210/pdb9VDR/pdb]: Crystal structure of voltage-gated sodium channel NavAb N49K/L176F mutant, 9VDS[https://doi.org/10.2210/pdb9VDS/pdb]: Crystal structure of voltage-gated sodium channel NavAb N49K/L176W mutant, 9VDT[https://doi.org/10.2210/pdb9VDT/pdb]: Crystal structure of voltage-gated sodium channel NavAb ΔC230/N49K mutant, 9VDU[https://doi.org/10.2210/pdb9VDU/pdb]: Crystal structure of voltage-gated sodium channel NavAb ΔC230/N49K/L176F mutant, 24JS[https://doi.org/10.2210/pdb24JS/pdb]: Crystal structure of voltage-gated sodium channel NavAb N49K/T206A mutant, 24JT[https://doi.org/10.2210/pdb24JT/pdb]: Crystal structure of voltage-gated sodium channel NavAb ΔC230/N49K/T206A mutant, 24JSU[https://doi.org/10.2210/pdb24JU/pdb]: Crystal structure of voltage-gated sodium channel NavAb ΔC230/N49K/L176F/T206A mutant. The structure of the NavAb N49K mutant for the initial model of molecular replacement was available in the Protein Data Bank under accession code 5YUC. The electrophysiologic data that support this study are available from the corresponding author, K.I., upon reasonable request.

## Acknowledgments

This work was funded by NIH grant 1R01GM142816 (SW). The synchrotron radiation experiments were performed at BL41XU and BL45XU in SPring-8 with the approval of the Japan Synchrotron Radiation Research Institute (JASRI) (Proposal numbers 2013B1178, 2015B1042, 2016B2721, 2017B2735, and 2018B2710). We thank the beamline staff for their excellent facilities and support. This research was partially supported by Platform Project for Supporting Drug Discovery and Life Science Research (Basis for Supporting Innovative Drug Discovery and Life Science Research [BINDS]) from AMED under Grant Numbers JP21am0101070, JP22ama121001 and JP23ama121001. This work was supported by Grants-in-Aid for Scientific Research (17K17795, 21K19332, 20K09193, and 24K02168), SEI Group CSR Foundation, Takeda Science Foundation, and the Institute for Fermentation (KI).

## Author Contributions

SW and KI conceived the studies. SW and SH designed and performed smFRET studies and analyzed smFRET data with the help of SA and JV. KI designed and performed electrophysiological and crystallographic studies and analyzed the resulting data. YM collected and analyzed electrophysiological data. SW and KI prepared the manuscript with editing input from other authors.

## Notes

### Competing Interest Statement

The authors have declared no competing interest.

### Summary of Updates

The revised manuscript included new crystal structures and analyses that uncovered the conformational dynamics underlying slow inactivation in the prokaryotic voltage-gated sodium channel NavAb and the critical residue pair coupling the conformational changes between the slow inactivation gate at the selectivity filter and the primary gate at the helix bundle crossing.

