## Extended Figure 1-6 for "The structural dynamics and molecular coupling in the slow inactivation of a prokaryotic voltage-gated sodium channel"

*Authors contributed to this work equally

[#]Corresponding to Katsumasa Irie and Shizhen Wang

**Extended Table 1. Data collection and refinement statistics.**

| | BG(N49K) | L176F | L176W | ΔC230 | ΔC230/ L176F |
|---|---|---|---|---|---|
| **PDB code** | 9VDQ | 9VDR | 9VDS | 9VDT | 9VDU |
| **Resolution range** | 41 - 3.1 (3.2-3.1) | 30 - 3.7 (3.8-3.7) | 30 - 3.7 (3.8-3.7) | 29.7- 3.5 (3.6-3.5) | 29.3 - 3.4 (3.5 - 3.4) |
| **Space group** | I 4 2 2 | I 4 2 2 | I 4 2 2 | I 4 2 2 | I 4 2 2 |
| **Unit cell** | 127.2 127.2 201.4 | 125.3 125.3 202.4 | 126.9 126.9 204.9 | 129.8 129.8 200.3 | 128.5 128.5 200.8 |
| **Unique reflections** | 15361 (1079) | 8934 (828) | 9266 (869) | 11127 (980) | 11918 (1085) |
| **Multiplicity** | 14.6 (15.0) | 77.7 (76.6) | 54.4 (55.4) | 26.1 (27.2) | 104.7 (103.9) |
| **Completeness (%)** | 96.15 (71.03) | 99.03 (94.85) | 99.30 (96.66) | 98.75 (90.49) | 99.15 (94.43) |
| **Mean I/sigma(I)** | 21.38 (1.95) | 31.17 (5.09) | 26.57 (4.83) | 27.92 (3.01) | 24.75 (3.44) |
| **Wilson B-factor** | 65.66 | 108.33 | 152.50 | 97.64 | 88.46 |
| **R-pim** | 0.0348 (0.4332) | 0.0122 (0.1726) | 0.0172 (0.4321) | 0.0168 (0.2434) | 0.0325 (0.3729) |
| **CC1/2** | 0.999 (0.886) | 1 (0.994) | 1 (0.986) | 0.999 (0.979) | 0.985 (0.932) |
| **Reflections used in refinement** | 14806 (1079) | 8880 (828) | 9236 (869) | 11025 (980) | 11854 (1085) |
| **R-work** | 0.241 (0.294) | 0.298 (0.294) | 0.313 (0.345) | 0.308 (0.272) | 0.254 (0.246) |
| **R-free** | 0.289 (0.357) | 0.327 (0.445) | 0.339 (0.413) | 0.344 (0.310) | 0.304 (0.270) |
| **Number of non-** | 2281 | 2044 | 1866 | 2225 | 2155 |

| | | | | | |
|---|---|---|---|---|---|
| hydrogen atoms | | | | | |
| macromolecules | 1870 | 1832 | 1798 | 1784 | 1795 |
| ligands | 401 | 207 | 68 | 436 | 355 |
| solvent | 10 | 5 | | 5 | 5 |
| Protein residues | 229 | 223 | 218 | 217 | 218 |
| RMS(bonds) | 0.012 | 0.004 | 0.003 | 0.003 | 0.010 |
| RMS(angles) | 1.32 | 0.71 | 0.48 | 0.76 | 1.27 |
| Ramachandran favored (%) | 99.12 | 99.09 | 98.13 | 100.00 | 99.53 |
| Ramachandran allowed (%) | 0.88 | 0.91 | 1.87 | 0.00 | 0.47 |
| Ramachandran outliers (%) | 0.00 | 0.00 | 0.00 | 0.00 | 0.00 |
| Rotamer outliers (%) | 0.96 | 0.00 | 0.00 | 0.00 | 0.00 |
| Clashscore | 15.07 | 9.39 | 6.25 | 9.88 | 13.98 |
| Average B-factor | 68.96 | 124.66 | 163.57 | 126.84 | 89.12 |

Statistics for the highest-resolution shell are shown in parentheses.

**Extended Table 2. Data collection and refinement statistics for T206A mutants.**

| | T206A | ΔC230/T206A | ΔC230/L176F/T206A |
|---|---|---|---|
| **PDB code** | 24JS | 24JT | 24JU |
| **Resolution range** | 44.3 - 3.5 (3.6 - 3.5) | 44.2 - 4.1 (4.2 - 4.1) | 45.3 - 3.7 (3.8 - 3.7) |
| **Space group** | I 4 2 2 | I 4 2 2 | I 4 2 2 |
| **Unit cell** | 127.0 127.0 203.4 | 128.3 128.3 202.4 | 128.0 128.0 205.2 |
| **Unique reflections** | 10874 (803) | 6934 (263) | 9451 (518) |
| **Multiplicity** | 90.9 (90.6) | 103.9 (81.2) | 100.9 (90.1) |
| **Completeness (%)** | 96.66 (74.98) | 89.27 (38.65) | 93.89 (56.49) |
| **Mean I/sigma(I)** | 21.15 (3.11) | 11.31 (1.04) | 14.51 (1.64) |
| **Wilson B-factor** | 68.82 | 178.88 | 79.01 |
| **R-pim** | 0.0708 (-2.826) | 0.0965 (0.5743) | 0.1595 (-0.4798) |
| **CC1/2** | 0.983 (0.957) | 0.998 (0.448) | 0.998 (0.877) |
| **Reflections used in refinement** | 10533 (803) | 6218 (264) | 8892 (518) |
| **R-work** | 0.280 (0.310) | 0.354 (0.434) | 0.279 (0.255) |
| **R-free** | 0.294 (0.292) | 0.375 (0.429) | 0.320 (0.377) |
| **Number of non-hydrogen atoms** | 1956 | 1779 | 2212 |
| **macromolecules** | 1797 | 1711 | 1778 |

| | | | |
|---|---|---|---|
| **ligands** | 159 | 68 | 434 |
| **Protein residues** | 222 | 211 | 227 |
| **RMS(bonds)** | 0.012 | 0.005 | 0.012 |
| **RMS(angles)** | 1.32 | 0.90 | 1.35 |
| **Ramachandran favored (%)** | 98.60 | 99.02 | 99.06 |
| **Ramachandran allowed (%)** | 1.40 | 0.98 | 0.94 |
| **Ramachandran outliers (%)** | 0.00 | 0.00 | 0.00 |
| **Rotamer outliers (%)** | 0.00 | 0.00 | 0.00 |
| **Clashscore** | 12.75 | 19.40 | 12.47 |
| **Average B-factor** | 68.20 | 194.81 | 87.98 |

Statistics for the highest-resolution shell are shown in parentheses.

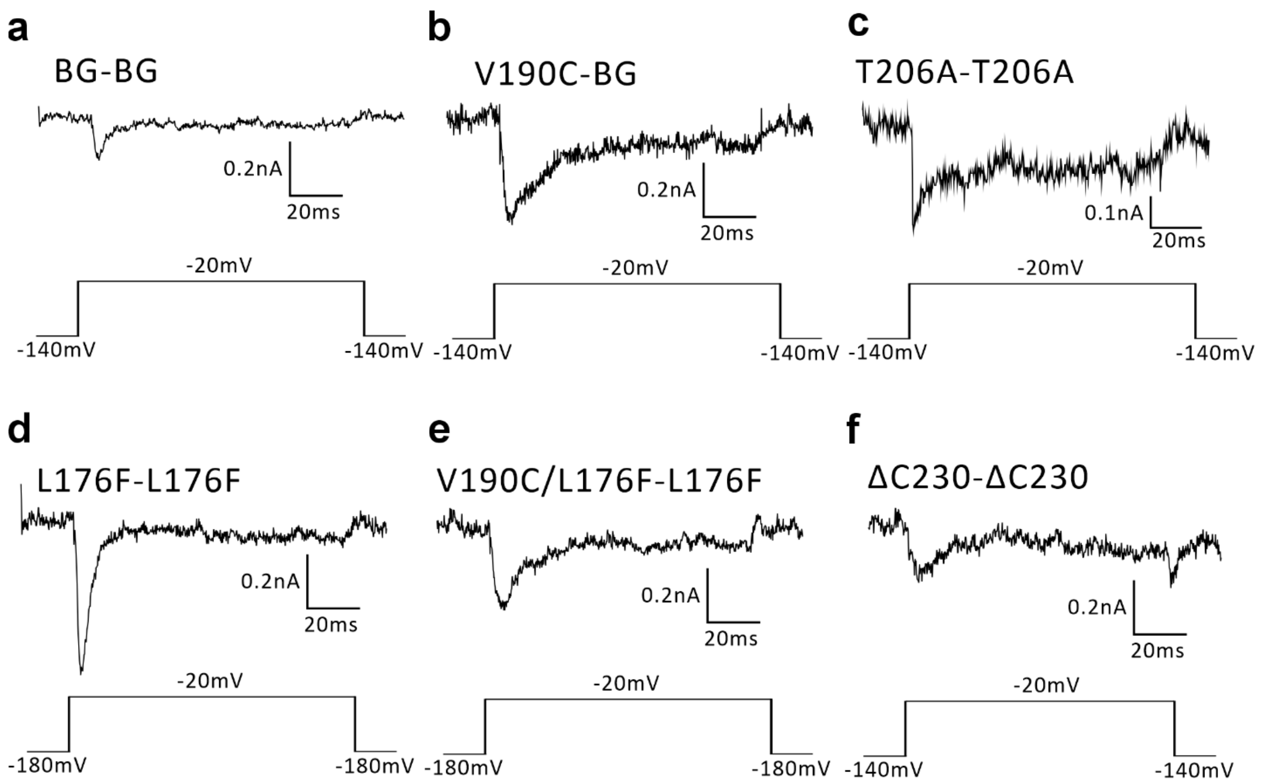


**Extended Figure 1. The channel activity of NavAb tandem dimers**

Representative trace of NavAb (**a**) BG dimer carrying only the N49K background mutation, (**b**) BG dimer with additional V190C in first subunit, (**c**) BG dimer with additional T206A mutation in both subunits, (**d**) BG dimer with additional L176F mutation in both subunits, (**e**) BG dimer with additional L176F mutation in both subunits and V190C in first subunit, and (**f**) BG dimer with C-terminal deletion ΔCHB in both subunits. All currents were generated by stimulation pulses of -20 mV.

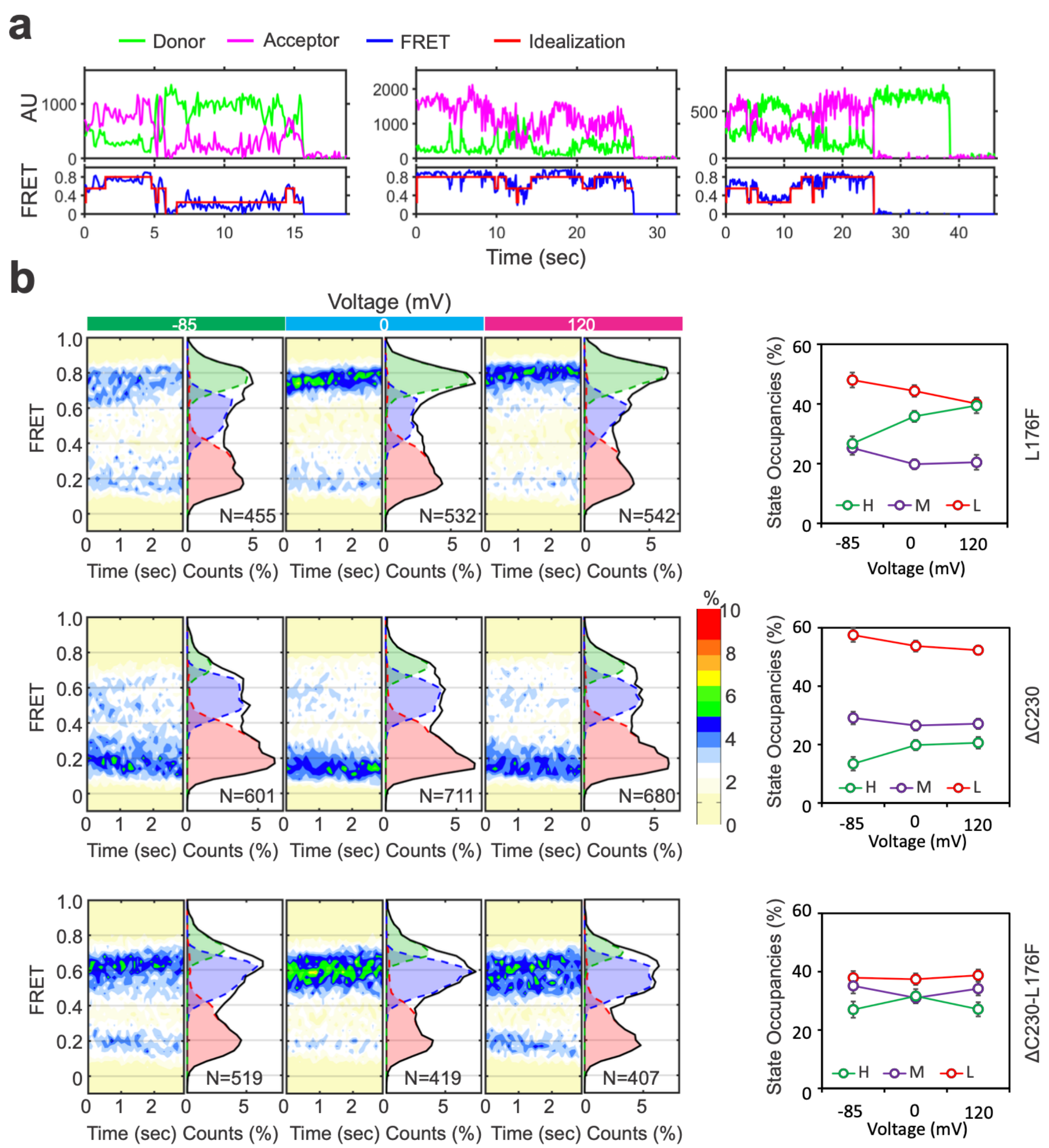


**Extended Figure 2. Voltage-dependent conformational changes in the selectivity filter of the NavAb channels carrying different mutations collected from the V190C labeling sites**

**a**. Representative smFRET traces exhibiting conformational transitions among 3 different states collected from V190C labeling sites; **b**. Histograms, contour maps and FRET state occupancies of smFRET data collected from either V190C labeling sites carrying either L176F, ΔC230 or L176F/ΔC230 mutations, under -85, 0 and 120 mV.

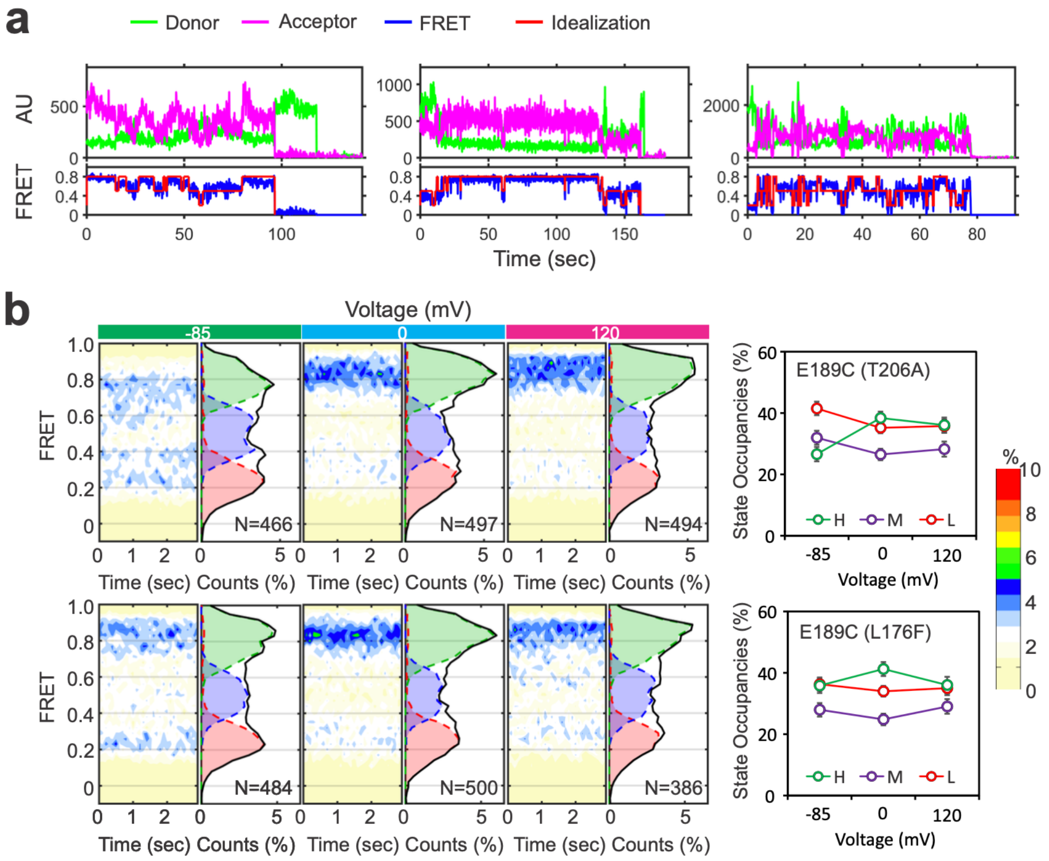


**Extended Figure 3. Voltage-dependent conformational changes in the selectivity filter of the NavAb channels carrying different mutations collected from the V189C labeling sites**

**a**. Representative smFRET traces exhibiting conformational transitions among 3 different states collected from V189C labeling sites; **b**. Histograms, contour maps and FRET state occupancies of smFRET data collected from either V189C labeling sites carrying either L176F and T206A mutations, under -85, 0 and 120 mV.

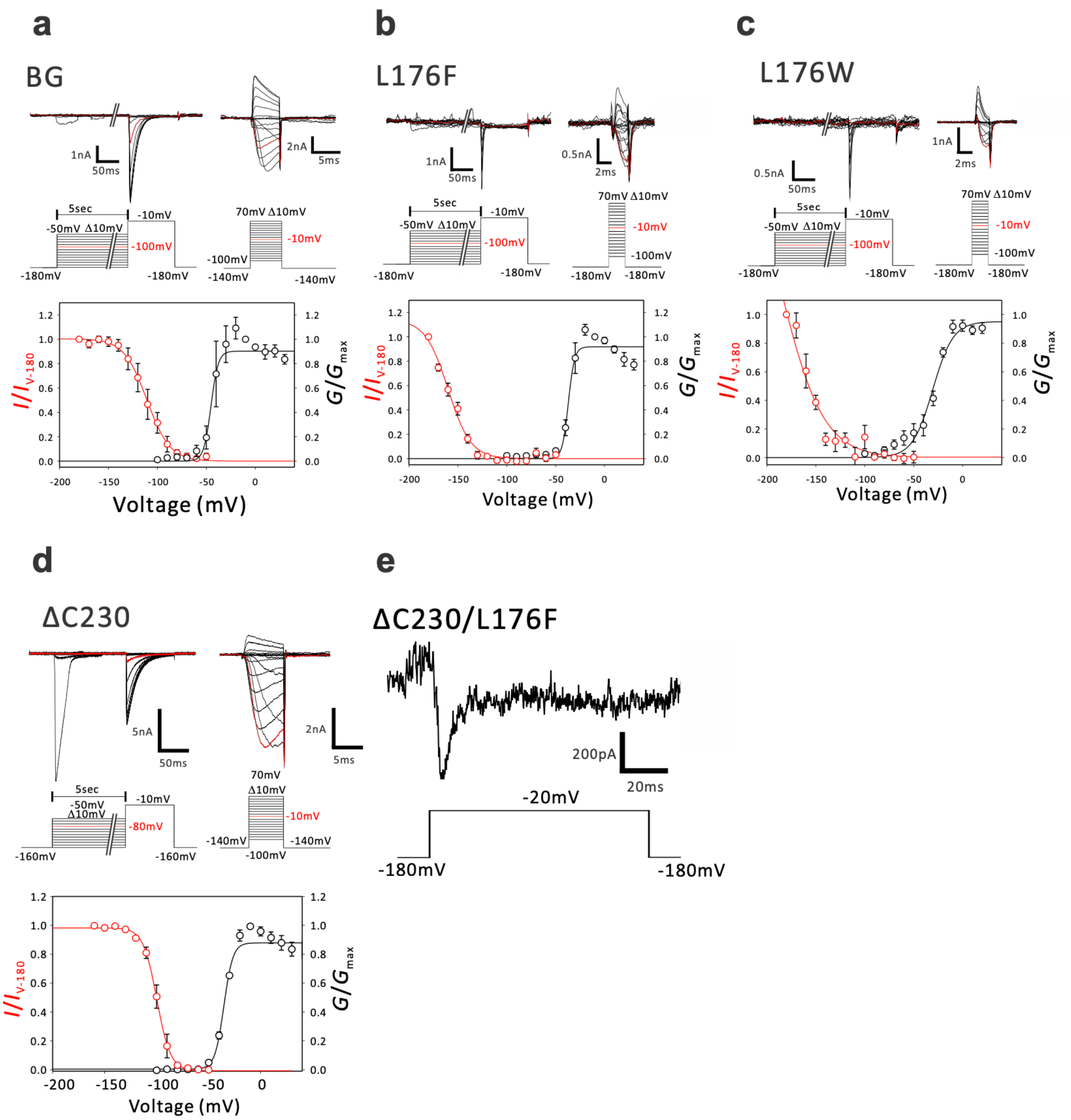

**Extended Figure 4. Voltage-dependent activation and inactivation of BG, L176F, L176W, ΔC230 and ΔC230/L176F mutants.**

**a-d**. Representative traces of deactivation tail currents and steady-state inactivation, and voltage-dependency of activation and inactivation of BG, L176F, L176W and ΔC230 mutants. **e**. Representative trace of the ΔC230/L176F current generated by stimulation pulses of -20 mV.

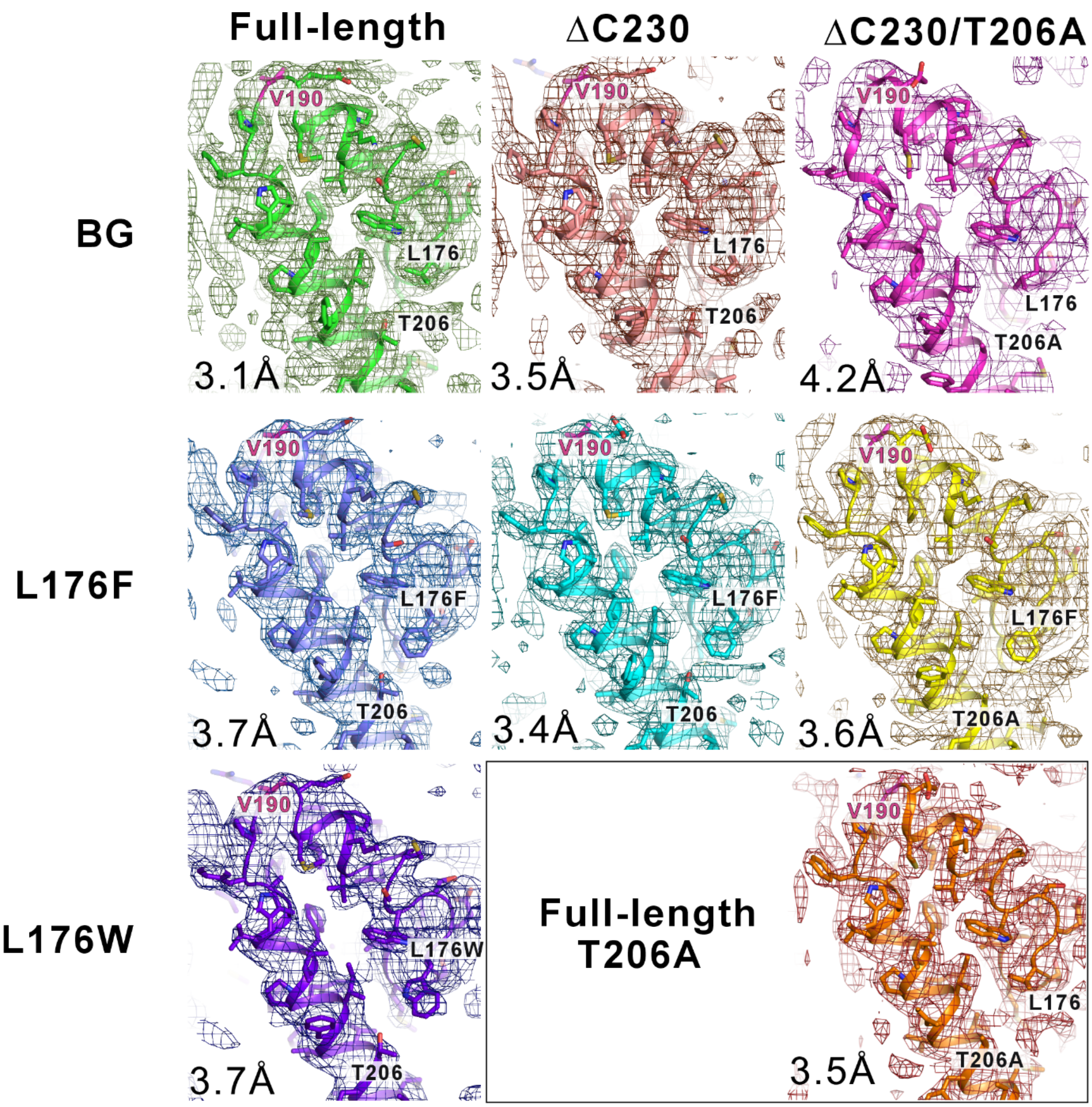


**Extended Figure 5. The electron density maps of the selectivity filters, P2 helices, and the extracellular side of the S6 helices in the NavAb mutants.**

2*Fo*-*Fc* electron density map with the amino acid residues surrounding the SF, P2 helix, and the extracellular side of the S6 helix. All mutants carry the N49K background mutation present in the BG mutant. Each mesh indicates the electron density maps contoured at 1σ. The structure resolutions are indicated in the bottom left of each panel.

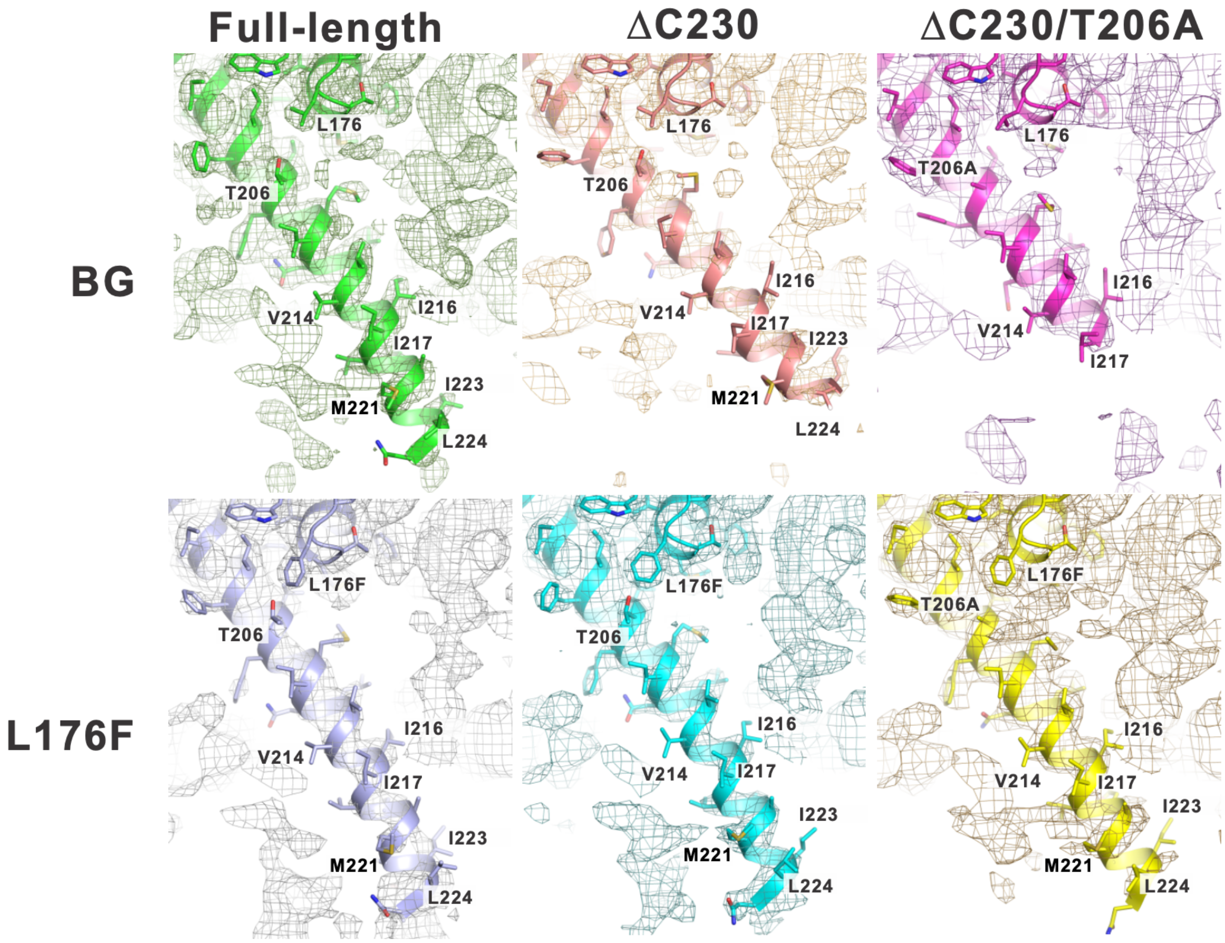


**Extended Figure 6. The omit map of the S6 helices in the NavAb channel carrying different mutations.**

An omitted *Fo* electron density map with the omitted amino acid residues of the S6 helix (from residue 203 to 224). All mutants carry the N49K background mutation present in the BG mutant. Each mesh indicates the electron density maps contoured at 0.8σ.
